# A Stealthy Player in Lipid Experiments? EDTA Binding to Phosphatidylcholine Membranes Probed by Simulations and Monolayer Experiments

**DOI:** 10.1101/2023.03.12.532294

**Authors:** Katarina Vazdar, Carmelo Tempra, Agnieszka Olżyńska, Denys Biriukov, Lukasz Cwiklik, Mario Vazdar

**Affiliations:** J. Heyrovský Institute of Physical Chemistry, Czech Academy of Sciences, Dolejškova 3, 18223 Prague, Czech Republic; Institute of Organic Chemistry and Biochemistry of the Czech Academy of Sciences, Flemingovo náměstí 542/2, 16000 Prague, Czech Republic; Central European Institute of Technology, Masaryk University, Kamenice 5, 625 00 Brno, Czech Republic; Department of Mathematics, Informatics and Cybernetics, University of Chemistry and Technology, Technická 5, 16628 Prague, Czech Republic

## Abstract

Ethylenediaminetetraacetic acid (EDTA) is frequently used in lipid experiments to remove redundant ions, such as Ca^2+^, from the sample solution. In this work, combining molecular dynamics (MD) simulations and Langmuir monolayer experiments, we show that on top of the expected Ca^2+^ depletion, EDTA anions themselves bind to phosphatidylcholine (PC) monolayers. This binding, originating from EDTA interaction with choline groups of PC lipids, leads to the adsorption of EDTA anions at the monolayer surface and concentrationdependent changes in surface pressure as measured by monolayer experiments and explained by MD simulations. This surprising observation emphasizes that lipid experiments carried out using EDTA-containing solutions, especially of high concentrations, must be interpreted very carefully due to potential interfering interactions of EDTA with lipids and other biomolecules involved in the experiment, e.g., cationic peptides, that may alter membrane-binding affinities of studied compounds.

**TOC Figure:** 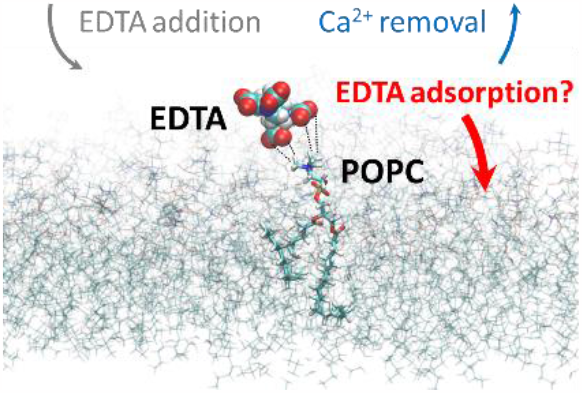

---

Ethylenediaminetetraacetic acid (EDTA) is a long-used synthetic aminopolycarboxylic acid prepared for the first time in 1935 and mainly known for its metal chelating properties.^1–3^ It is effectively utilized for the complexation of several multivalent cations but most frequently is applied for forming water-soluble complexes with iron (both Fe^2+^ and Fe^3+^)^4,5^ and calcium Ca^2+^ ions at neutral pH in solution.^6^ Due to its versatile metal chelating properties,^7^ it has found a staggering number of applications in industry, medicine, and household. EDTA is used for treating mercury and lead poisoning^8^, as a preservative in different drug formulations,^9,10^ food industry^11^, and in the cosmetic industry for stabilization of formulations during air exposure.^12^ Additionally, EDTA, as well as other metal chelators are known to induce permeabilization of the outer membrane in Gram-negative bacteria.^13^ It is speculated that EDTA actively removes metal ions from the bacterial membrane, resulting in the loss of lipopolysaccharides and proteins, leading to cell lysis.^14^ On the fun side, due to all of the listed various EDTA applications in academic and commercial applications, “strong additional evidence of the efficient use” of EDTA has also been reported in popular culture^15^.

In academic research, EDTA has an invaluable position as an efficient metal chelator for removing redundant ions in solution^16^ and biological membranes^17^ or inhibiting metal-dependent proteins.^18,19^ As such, it is almost always used as an additive in the preparation of a whole range of buffers in biological and biophysical investigations on cells and liposomal cell models, with the aim of total removal of leftover calcium Ca^2+^ ions in Milli-Q water (which are inevitably present in the low nM concentration in our experimental conditions) as well as its sequestration from the biological membranes where they easily bind.^20^ Surprisingly, there are very few studies that aimed to test whether EDTA itself binds to lipids and what could be possible consequences of such interaction. For instance, EDTA was noted to have an enhancing effect on the action activity of antiglaucoma drugs by increasing the permeability of the corneal membrane.^21^. Galla et al. have reported the fluidization and expansion effect of EDTA in higher concentrations on the phase behavior of dipalmitoylphosphatidylcholine (DPPC) monolayers, brought upon by the electrostatic interaction of negatively charged carboxylic groups of EDTA with the positively charged headgroup of DPPC.^22^ Using AFM, they have shown that intercalation of EDTA in the DPPC monolayer induces a membrane curvature, which size and magnitude depend on the length of exposure to EDTA. However, the molecular picture of the interaction has been only qualitatively described using simple molecular mechanics and semiempirical PM3 calculations on solvent-free models, thereby entirely disregarding the dynamical component of the interaction, which might be relevant for the described membrane curvature changes.^22^ Essentially, to the best of our knowledge, EDTA action on membranes has not been carefully considered yet at the molecular level, and its sequestration of Ca^+2^ ions from lipid membranes is very often taken for granted without a sufficient understanding of whether the addition of EDTA to biological samples has any other effect on corresponding experiments. Considering that only a minimal concentration of Ca^2+^ is always present in Milli-Q water (in a low nM concentration range) while EDTA is added in much higher mM concentration to lipid systems (ranging from 0.1 mM^23,24^ to even 5 mM in some buffers used in cell biological experiments for membrane protein extraction^25,26^), an obvious possible interaction of the significant excess of EDTA anions with lipid membranes leading to their adsorption and consequent implications has surprisingly never been studied. In this work, we tackle this open question and systematically investigate the adsorption of EDTA anions (whose distribution depending on the pH of the solution is shown in Figure 1) to 1-palmitoyl-2-oleoylphosphatidylcholine (POPC) monolayers as the simplest model of lipid membranes combining the custom Langmuir-trough monolayer experiments with computer simulations.

**Figure 1.**
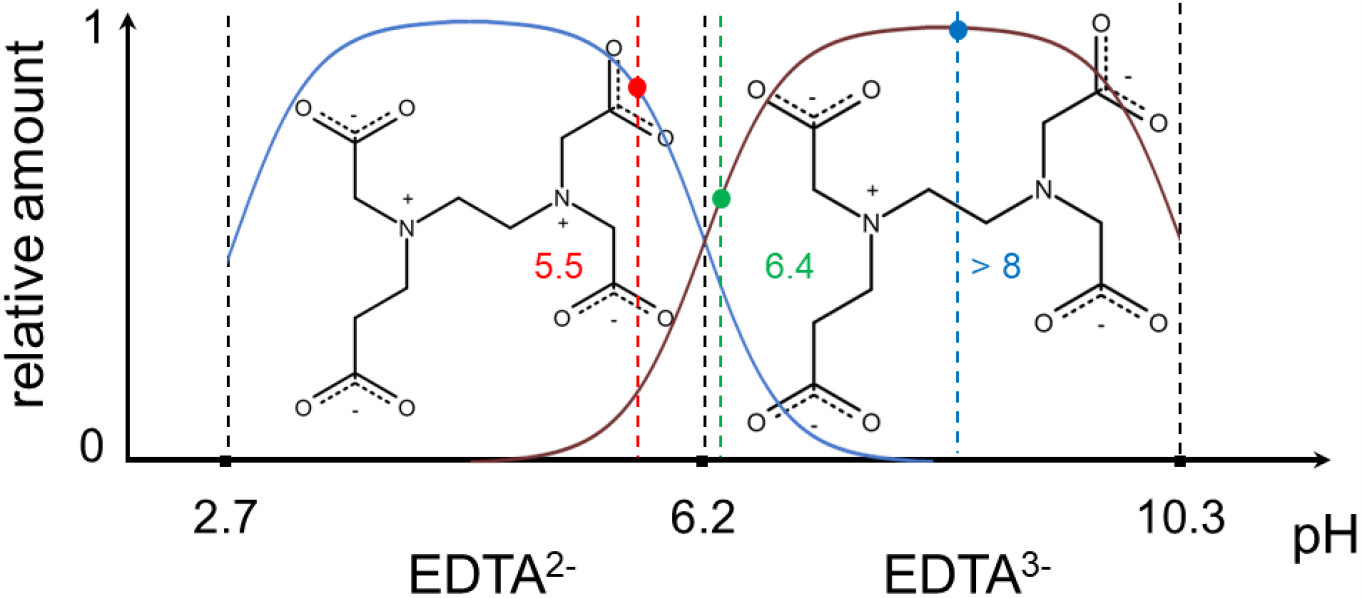
A relative amount of EDTA^2-^ (blue line) and EDTA^3-^ (brown line) in water according to the Henderson-Hasselbalch equation^27^ as a function of corresponding EDTA p*K*_a_ values. For low 50 nM EDTA experiments, the measured pH of the solution is 5.5, and EDTA is present as EDTA^2-^ in ca. 80% and EDTA^3-^ ca. 20% (red dot). At 50 μM EDTA concentration, pH is 6.4, and the ratio between EDTA^2-^ and EDTA^3-^ is similar (green dot). At higher EDTA concentrations (1 mM, 3 mM, 15 mM, and 150 mM), the measured pH values are slightly larger than 8, and almost all EDTA ions in the solution are in the EDTA^3-^ form (blue dot).

First, we performed MD simulations of 1-palmitoyl-2-oleoylphosphatidylcholine (POPC) monolayers interacting with EDTA^2-^ and EDTA^3-^ anions at different EDTA / Ca^2+^/POPC ratios (exact composition of the systems is presented in Table S1), and the results are presented in Figure 2 (Simulation Details are explained in detail in the SI). Note that we examine the adsorption of EDTA in two possible protonation states according to corresponding p*K*_a_ values and pH, see Figure 1. The analysis of number density profiles in 1 EDTA^2-^ / 10 POPC and 1 EDTA^3-^ / 10 POPC systems (upper left and right panels, respectively) shows that despite the high negative charge of EDTA anions, both EDTA^2-^ and EDTA^3-^ have slightly pronounced adsorption peaks at around 1 nm from phosphate POPC atoms. This weak adsorption is present for both EDTA anions and comparable in strength (if not even stronger) to adsorption of only singly negatively charged ions such as Cl^−^.^28^ Interestingly, the difference in charge between EDTA^2-^ and EDTA^3-^ does not contribute to the strength of adsorption, because carboxyl groups of both EDTA anions similarly interact with positively charged POPC choline groups. The density profile of Na^+^ counterions (green color) shows only a small amount close to the membrane surface. However, note that a recent study combining MD simulations, vibrational sum frequency generation, and Langmuir-trough experiments has shown that the presence of Na^+^ cations at DPPC monolayers does not disturb the membrane even at high mM concentrations,^29^ implying that any changes in monolayer structure are not induced by sodium and should be attributed to other ions and solutes present in the solution.

**Figure 2.**
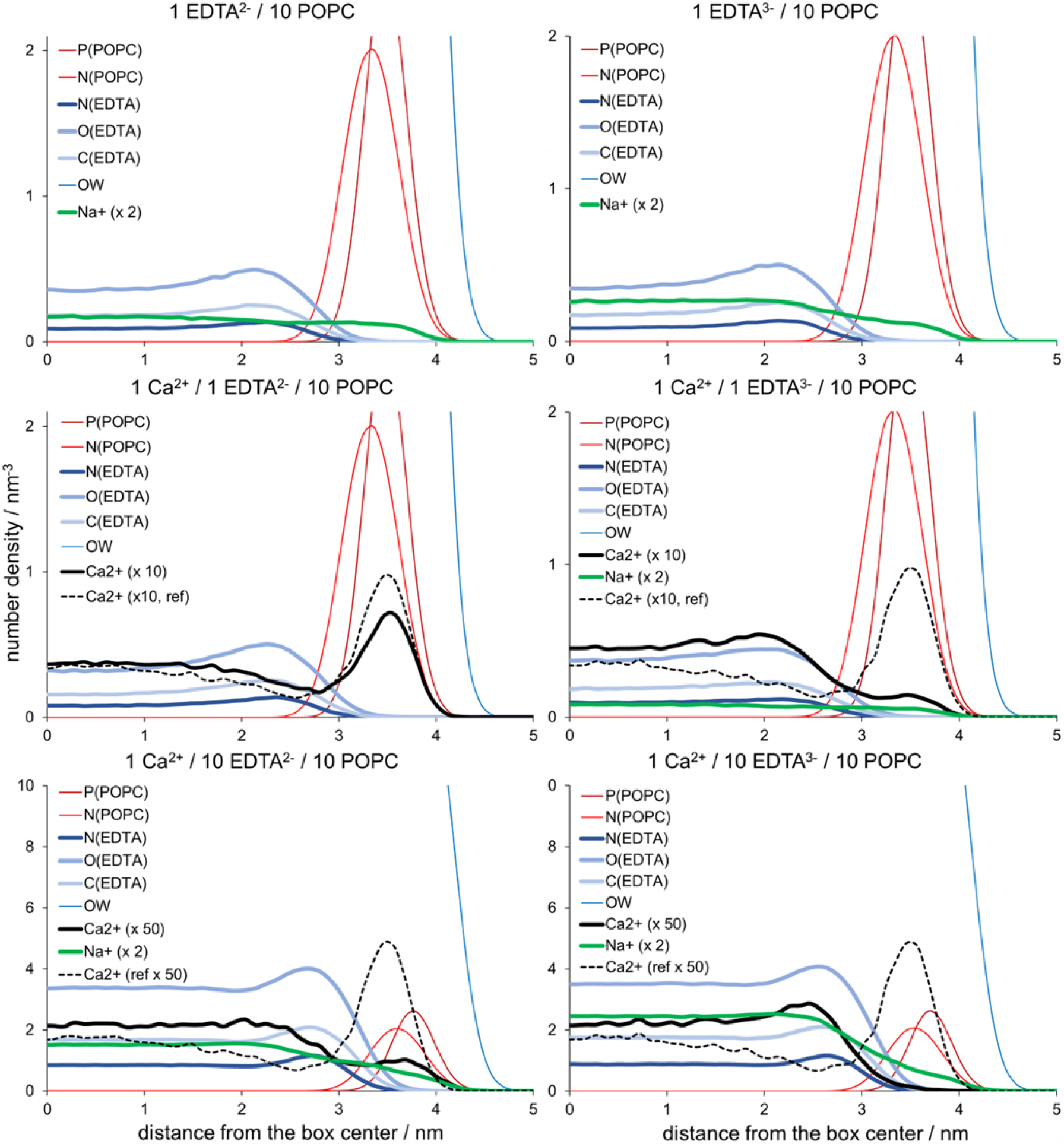
Number density profiles for monolayer phosphate atoms P(POPC), choline nitrogen atoms N(POPC), all nitrogen (N(EDTA)), oxygen (O(EDTA)), and carbon atoms of EDTA (C(EDTA)), Na^+^ and Ca^2+^ cations, and water oxygen atoms from MD simulations with different ratios of EDTA / Ca^2+^ / POPC. The number density profiles for EDTA atoms and cations are shown in thick lines. The number density of calcium is scaled up by a factor of 10 or 50 (depending on the EDTA concentration) to highlight the effect of its sequestration. Similarly, the number density of sodium is scaled up by a factor of 2. The reference number density profile for calcium from EDTA-free systems is shown in a dashed line.

In 1 Ca^2+^/1 EDTA^2-^/10 POPC system (middle left panel, Figure 2), which is set up to mimic the experiments at low nM EDTA concentrations at pH around 5.5 (Figure 1), we observe that in addition to adsorption of EDTA^2-^, which exhibits an almost identical adsorption pattern as in the reference system (left upper panel), Ca^2+^ ions are still relatively abundant at the POPC monolayer. This observation is evidenced by a higher number density of Ca^2+^ in the headgroup region vs. bulk Ca^2+^ concentration in our MD simulations, but still in a smaller amount than in EDTA-free simulations containing only Ca^2+^ ions (dashed lines). Therefore, MD simulations indicate only a partial sequestration of Ca^2+^. Upon extra addition of EDTA^2-^ to the system (1 Ca^2+^/10 EDTA^2-^/10 POPC, bottom left panel), the number density of Ca^2+^ shows its further removal from the monolayer surface. However, we should note that such EDTA^2-^ to calcium ratio is not experimentally observed since adding EDTA increases the pH and, as a result, also increases the EDTA^3-^ concentration at the expense of EDTA^2-^ anions (Figure 1).

Therefore, we also modeled EDTA^3-^-containing systems which more closely correspond to the experiments with higher EDTA concentrations where the pH is around 6.4 (Figure 1). The overall adsorption of EDTA^3-^ is similar in 1 Ca^2+^/1 EDTA^3-^/10 POPC vs. 1 Ca^2+^/1 EDTA^2-^ /10 POPC (Figure 2, middle panels), with one important exception – the sequestration of Ca^2+^ is more efficient in EDTA^3-^ system compared to analogous EDTA^2-^ and reference EDTA-free systems. Finally, in 1 Ca^2+^/10 EDTA^3-^/10 POPC system, the sequestration of Ca^2+^ from the monolayer is complete (Figure 2, bottom right panel), indicating more efficient Ca^2+^ depletion with increasing EDTA^3-^ concentration, which is intuitively expected due to the stronger electrostatic Ca^2+^-EDTA^3-^ interaction vs. Ca^2+^-EDTA^2-^ interaction. Altogether, our conclusions drawn from MD simulations perfectly resemble the anticipated function of EDTA, yet indicating that in all cases, some amount of EDTA remains bound to the lipid monolayer.

For the first experimental measurements (for Experimental Details see the SI), we decided to check the effect of low 50 nm EDTA concentrations at pH = 5.5, where the concentration of EDTA is comparable to the Ca^2+^ concentration in Milli-Q water (used in our experiments) as modeled in 1 Ca^2+^/1 EDTA^2-^/10 POPC system. Using a custom-made micro-well with the possibility of substance injection into the subphase and equipped with a surface tensiometer (see Methods for details), we measured how the addition of 50 nM of EDTA affects the surface pressure of the POPC monolayer. We observed only a slight decrease in the surface pressure with time; the POPC monolayer stabilized with surface pressure ca. 0.5 mN m^−1^ lower than before the EDTA addition (Figure 3, top left panel). The drop of pressure is attributed to only a partial removal of Ca^2+^ from the monolayer headgroup region in agreement with a minor decrease in the number density of Ca^2+^ in 1 Ca^2+^/1 EDTA^2-^/10 POPC system vs. reference EDTA-free system (Figure 2, left middle panel).

**Figure 3.**
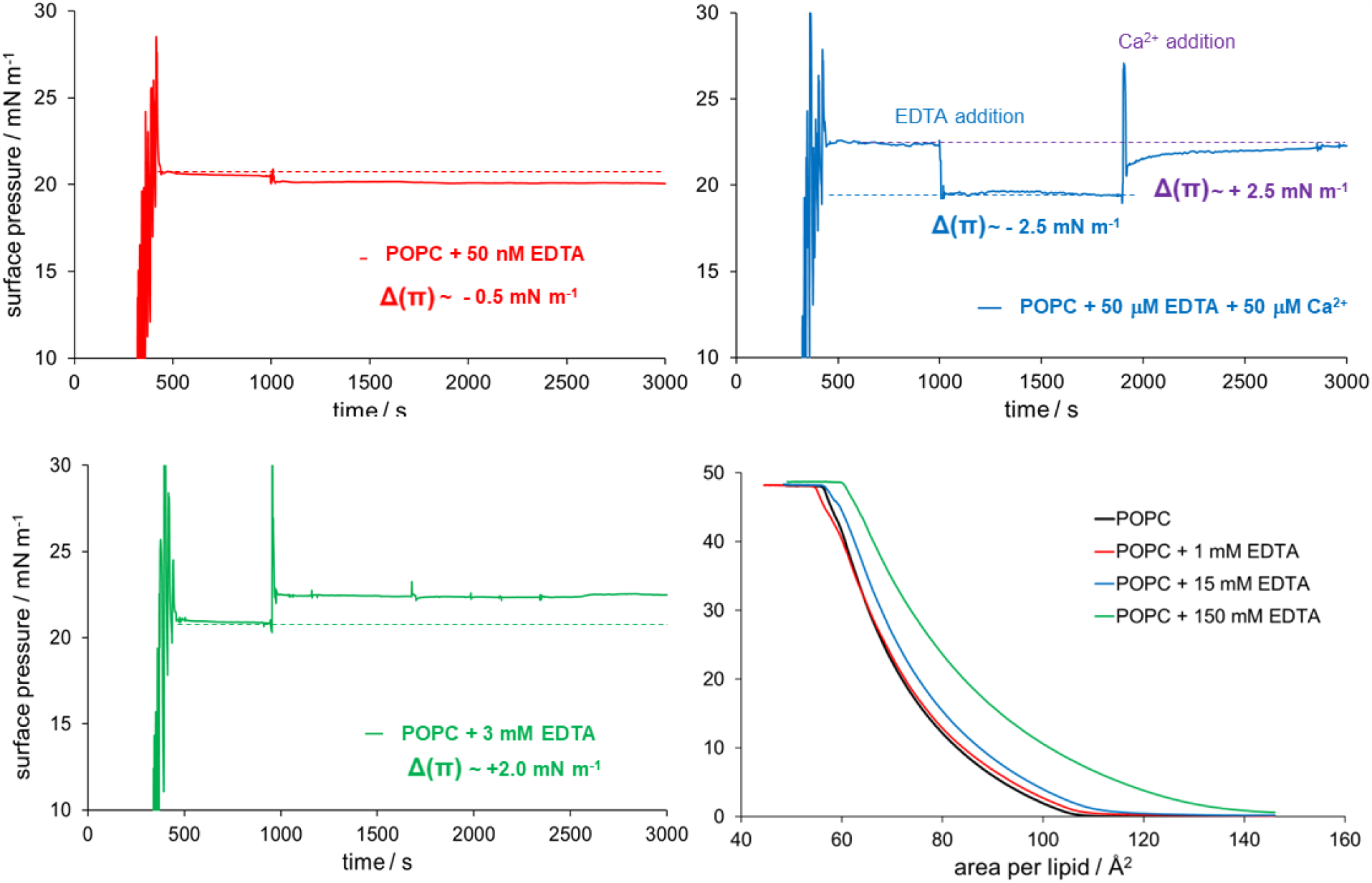
Kinetics of POPC monolayer before and after addition of EDTA (top left and bottom left panel) and EDTA with subsequent CaCl_2_. The EDTA concentrations in the solution are 50 nM at pH = 5.5 (red, top left panel), 50 μM at pH = 6.4 (blue, top right panel), and 3 mM at pH = 8 (green, bottom left panel). Langmuir compression isotherms of POPC in the presence of EDTA at different concentrations (multicolor, bottom right panel).

Since the removal of Ca^2+^ is incomplete under these experimental conditions, we performed additional experiments at 50 μM EDTA concentration, where the measured pH is 6.4. At this EDTA concentration, the amount of EDTA^3-^ increased vs. EDTA^2-^, now them being roughly in the same amount in the solution (Figure 1). An observed drop of the surface pressure after EDTA addition significantly increased from ca. 0.5 to 2.5 mN m^−1^, which agrees with 1 Ca^2+^/1 EDTA^3-^/10 POPC and 1 Ca^2+^/10 EDTA^3-^/10 POPC density profiles (Figure 2, middle right and middle bottom panel), indicating that Ca^2+^ is more efficiently removed from the POPC surface by EDTA^3-^, especially in the case of 1 Ca^2+^/10 EDTA^3-^/10 POPC system where Ca^2+^ is completely depleted from the lipid monolayer (Figure 2, bottom right panel). Moreover, the subsequent addition of 50 μM Ca^2+^ (by corresponding addition of CaCl_2_ in the subphase) to the same experimental system showed an increase in surface pressure back to the level before the addition of EDTA (Figure 3, top right panel), thus again confirming that EDTA indeed removes Ca^2+^ from the membrane. Finally, we performed adsorption kinetics measurements at 3 mM EDTA concentrations (where pH is slightly larger than 8), which show the increase of surface pressure upon EDTA addition by ca. 2 mN m^−1^ (Figure 3, left bottom panel) due to the elevated effect of EDTA adsorption at higher concentrations, which cancels the effect of Ca^2+^ removal. These results are also confirmed by the independent experiments with different monolayer surface pressure sensors shown in Figure S1.

The Langmuir compression isotherms of POPC monolayers were measured on subphases of Milli-Q water (used as a reference) at 1 mM, 15 mM, and 150 mM of EDTA, see Figure 3, bottom right panel. The measured experimental pH values are slightly larger than 8, indicating that EDTA^3-^ anions are dominantly present in the solution (Figure 1). The interaction of EDTA^3-^ anions with the POPC monolayer is clearly visible at 15 mM and 150 mM EDTA concentrations. The isotherms are shifted horizontally (for area per lipid) and vertically (for surface pressure), and the effect increases with the concentration of added EDTA. The collected isotherms indicate that EDTA interacts and accumulates at the POPC monolayer, thereby increasing the surface pressure for the whole range of areas per lipid. Interestingly, for all EDTA concentrations, the measured isotherms coincide with the isotherm of Milli-Q water in the region of monolayer collapse at low area per lipid values (around 60 Å^2^), suggesting that a certain amount of EDTA remains trapped in the POPC monolayer until monolayer breakup.

Since the concentration of Ca^2+^ in Langmuir-trough experiments is by many orders of magnitude smaller than concentrations of EDTA^3-^ anions (nM and mM range, respectively), the best comparison with the MD simulation results can be made for Ca^2+^-free systems, in particular for the 1 EDTA^3-^/ 10 POPC system where EDTA^3-^ anions are most abundant species at experimental pH = 8. As indicated in the MD simulations section, the adsorption of EDTA^3-^ anions at the POPC monolayer is clearly observed for these conditions (Figure 2, upper right panel). At 1 mM concentration of EDTA, the isotherm is very similar to the referent POPC, and it is not completely clear whether 1 mM EDTA indeed leads to the observed shift in the isotherm and subsequent increase of surface pressure. However, we showed in adsorption kinetics measurements that at lower EDTA concentrations (50 nm and 50 μM), the surface pressure decreases due to at least partial Ca^2+^ sequestration from the membrane, whereas adding 3 mM EDTA leads in contrast to an increase in the surface pressure. Therefore, at the EDTA concentration of 1 mM, which shows only a small difference compared to the measurements on pure POPC, it is fair to assume that the opposite directions of the corresponding trends result in minimal (if any) changes in the surface pressure (Figure 3). In any case, we should stress that EDTA is still adsorbed at the POPC monolayer as shown in the MD simulations even with lower EDTA content (Figure 2).

From the MD simulation data and experimental surface pressure monolayer experiments, we can conclude the following. First, using MD simulations, we demonstrated that calcium sequestration is induced by both EDTA^2-^ and EDTA^3-^ anions already at low EDTA concentrations (Figure 2), which agrees with the experimental adsorption kinetics measurements. However, the fact that the Ca^2+^ sequestration at 50 nM and pH = 5.5 is only partial (Figure 2) suggests that higher concentrations of EDTA should be used for its complete removal. Indeed, in systems with higher EDTA concentrations of 50 μM (corresponding to pH=6.4, where a more significant amount of EDTA^3-^ is present in solution), MD simulations and experiments predict a more efficient removal of Ca^2+^ from the membrane (Figures 2 and 3). Moreover, the increase of EDTA concentration to 3 mM in adsorption kinetics measurements shows the increase of surface pressure induced by EDTA adsorption. Therefore, using 0.1 mM (or higher) EDTA concentrations in typical biophysical experiments is justified for successful Ca^2+^ removal from the lipid membranes.

Second, and far more intriguingly, we see that in MD simulations, both EDTA^2-^ and EDTA^3-^ anions adsorb to the POPC monolayer at all investigated EDTA concentrations, including the reference system without Ca^2+^. This observation is supported by a large surface pressure increase in kinetic and isotherm experimental measurements with high mM EDTA concentrations (Figure 3, bottom panels), in agreement with the increased number density of EDTA anions in corresponding MD simulations (Figure 2). These findings imply an additional electrostatic effect of EDTA weakly bound to the membrane. To check for that effect, we calculated the total electrostatic potential of the reference POPC system without EDTA and compared it with the 1 Ca^2+^/ 10 EDTA^3-^/ 10 POPC system (Figure S2). We see that in the EDTA-containing system, a small minimum of ca. 50 mV appears at around 3.5 nm away from the box center (red line), i.e., at the position of the number density profiles maxima of EDTA atom groups (Figure 2). This observation implies that the interaction of other positively charged species with the membrane might be partially screened by negatively charged EDTA, thus inhibiting the interaction with the POPC headgroups themselves.

In conclusion, we presented that the addition of EDTA in 0.1 mM concentrations is justified in biophysical and biological experiments since lower concentrations of EDTA do not lead to complete Ca^2+^ removal from the lipid membranes, as confirmed by both MD simulations and adsorption kinetics measurements. However, with the observed sequestration effect, an additional stealthy action of EDTA is also detected – its adsorption to POPC monolayers at all investigated EDTA concentrations. This behavior is evident from MD simulations but especially in kinetic and isotherm measurements showing an increase of surface pressure at the POPC monolayer with mM concentrations of EDTA. Moreover, given corresponding charge-screening effects induced by EDTA, its adsorption may influence the binding of other positively charged species (such as positively charged cell-penetrating peptides)^30,31^ especially when they are present in similarly low mM concentrations like 0.1 mM EDTA often used in biophysical experiments. One of the prominent examples where EDTA action might play a critical role is found in the lack of the translocation of polyarginine cell-penetrating peptides across large POPC unilamellar vesicles (LUVs) in EDTA-containing experiments^24^ vs. their facile penetration across giant unilamellar POPC vesicles (GUVs) in EDTA-free systems.^32,33^ Moreover, the energetics of peptide binding to POPC, known to be dependent also on the ionic strength of the solution,^34^ could change significantly when EDTA is present in the system, and the results of experiments involving EDTA should be taken with great care.

## Author Contribution

K. V., L. C., and M. V. conceptualized the study. C. T. and D. B. performed molecular dynamics simulations, whereas K. V. and A. O. performed the experimental part of the work. All authors interpreted the data, prepared the figures and M. V. wrote the first version of the manuscript. All authors revised and agreed on its final version.

## Data Availability

The data that support the findings of this study are available from the corresponding author upon reasonable request.

## Supporting Information

The Supporting Information is available free of charge at https://pubs.acs.org/doi/xxx.

- Composition of studied systems in MD simulations.
- Details of Simulation and Experimental Methods.
- Kinetics of POPC monolayer surface pressure changes.
- Calculated electrostatic potential in the studied systems.
- An example snapshot from EDTA-containing MD simulations.
- Simulation model of EDTA anions.
- Atomic partial charges of EDTA^2-^ and EDTA^3-^ anions.

## Acknowledgments

This work was supported by the Czech Science Foundation (grant no. 21-19854S) and the project “National Institute of Virology and Bacteriology (Program EXCELES, ID Project No. LX22NPO5103) - Funded by the European Union - Next Generation EU”.

