## Supporting Information for "A Stealthy Player in Lipid Experiments? EDTA Binding to Phosphatidylcholine Membranes Probed by Simulations and Monolayer Experiments"

Page 2. Composition of studied systems in MD simulations.

Page 3. Details of Simulation and Experimental Methods

Page 5. Kinetics of POPC monolayer surface pressure changes.

Page 6. Calculated electrostatic potential in the studied systems.

Page 7. An example snapshot from EDTA-containing MD simulations.

Page 8. Simulation model of EDTA anions.

Page 9. Atomic partial charges of $EDTA^{2-}$ and $EDTA^{3-}$ anions.

| **Table S1.** Composition of all studied systems in molecular dynamics simulations. | | | | | | | |
|---|---|---|---|---|---|---|---|
| Name | POPC | $EDTA^{2-}$ | $EDTA^{3-}$ | $Na^{+}$ | $Ca^{2+}$ | $Cl^{-}$ | Water |
| 1 $EDTA^{2-}$/10 POPC | 512 | 51 | 0 | 102 | 0 | 0 | 41380 |
| 1 $EDTA^{3-}$/10 POPC | 512 | 0 | 51 | 153 | 0 | 0 | 41360 |
| 1 $Ca^{2+}$/0 $EDTA^{2-}$/10 POPC | 512 | 0 | 0 | 0 | 51 | 102 | 41380 |
| 1 $Ca^{2+}$/1 $EDTA^{2-}$/10 POPC | 512 | 51 | 0 | 0 | 51 | 0 | 41380 |
| 1 $Ca^{2+}$/10 $EDTA^{2-}$/10 POPC | 512 | 512 | 0 | 922 | 51 | 0 | 40458 |
| 1 $Ca^{2+}$/1 $EDTA^{3-}$/10 POPC | 512 | 0 | 51 | 51 | 51 | 0 | 41357 |
| 1 $Ca^{2+}$/10 $EDTA^{3-}$/10 POPC | 512 | 0 | 512 | 1434 | 51 | 0 | 40248 |

**Methods**

**Simulation Details.** The initial POPC monolayer structure was taken from the publicly available data on the Zenodo server (DOI: 10.5281/zenodo.838633).[1] This structure contains a slab of water placed in the center of the box and two POPC monolayers (256 lipids each) located at the interfaces with a ~12 nm-thick vacuum (Figure S3). This system was used to prepare all other simulation setups by adding ions, including EDTA in different protonation forms. The complete composition of the simulated systems is summarized in Table S1. The obtained systems were energy minimized, and then 1 µs long production runs were carried out (only 500 ns for the reference system with $Ca^{2+}$ and no EDTA). The first 100 ns were considered as equilibration and disregarded from the analysis. All simulations were run using an NVT ensemble with an area per lipid of POPC of roughly 70 $Å^2$ (0.70 $nm^2$). The force field used for POPC and ions is CHARMM36,[2] while water was modeled using the four-point OPC model.[3] This combination of force fields was selected based on the recent work discussing the accuracy of lipid monolayer simulations.[4] The model for EDTA anions was built using Ligand Reader & Modeler module[5] in CHARMM-GUI[6], (Figure S4) and corresponding partial charges derived using CGenFF[7] are given in Table S2. The equation of motion was solved using 2 fs timestep and leap-frog algorithm, with an updating frequency of 20 steps. The smooth particle mesh Ewald with a cut-off of 1.2 nm was used to treat electrostatic interactions.[8] Van der Walls interactions were treated using a cut-off of 1.2 nm, with the forces smoothly attenuated to zero between 1.0 and 1.2 nm. The temperature was kept constant at 298 K using the Nose-Hoover thermostat with three coupling groups (POPC, EDTA anions, and remaining solvent). All covalent bonds involving hydrogens were constrained using the P-LINCS algorithm.[9] All simulations were performed using Gromacs, versions 2021 and 2022.[10]

**Experimental Details.** Measurements of monolayer surface pressure kinetics (adsorption kinetics) were performed with an in-house built micro-well (round shape interface of 7 $cm^2$, 5 mL subphase volume, perforation present for injection of solutions directly into subphase). The system was equipped either with an ultra-sensitive surface pressure sensor (Kibron) with the DyneProbe or NIMA surface tensiometer. 15 µL of 0.1 mM solution of POPC (purchased from Avanti Polar Lipids, Alabaster, AL) in chloroform were spread by deposition of small droplets with a Hamilton microsyringe over 4 mL of Milli-Q water (Millipore, USA, 18.2 MΩcm, pH 5.5, the concentration of $Ca^{2+}$ was in low nM range) to achieve a surface pressure of 20 mN $m^{-1}$. The surface pressure change was monitored for 30 mins needed for chloroform evaporation and monolayer stabilization. Once the surface pressure of POPC monolayer stabilized, EDTA and $CaCl_2$ in appropriate amounts (corresponding to concentrations of 50 nM, 50 μM, and 3 mM of EDTA, and 50 μM EDTA + 50 μM $CaCl_2$, respectively) were administered by Hamilton microsyringe to the subphase below the monolayer, and the pressure was monitored for further 30 minutes. Measurements were performed at room temperature. To slow down subphase evaporation and protect the film from dust and additional disruptions, an acrylic cover box over the setup was used.

Langmuir monolayer surface pressure-molecular area (π-A) compression isotherms were measured with a commercially available MicroTrough setup (59 mm × 209 mm) (μtrough XS, Kibron; Helsinki, Finland). The system was equipped with an ultra-sensitive surface pressure sensor (KBN 315; Kibron) with the DyneProbe. 12.5 μL of 1 mM solution of POPC in chloroform were spread by deposition of small droplets with a Hamilton microsyringe over 25 mL of Milli-Q water or solutions of EDTA of corresponding concentrations. The π-A isotherms were collected during the symmetrical movement of two barriers controlled by software (FilmWare) provided by the equipment manufacturer. The compression speed was 10 mm/min (i.e., 3.92 $Å^2$/chain/min for POPC monolayer). Measurements were done at 25.0 °C, controlled with a temperature control plate (connected to a water-circulating thermostat; ±0.5 °C accuracy) placed under the trough. To slow down subphase evaporation and protect from dust and additional surface disruptions, an acrylic cover box over the trough was used. Before each measurement, the lipid film was left uncovered for 3 min to allow chloroform to evaporate and then for 5 min covered with the acrylic box to enable the temperature to equilibrate.

150 mM solution of EDTA was prepared by dissolution of solid EDTA (Sigma-Aldrich; St. Louis, MO) and NaOH (Sigma-Aldrich; St. Louis, MO) in Milli-Q water, and the final pH was set to 8.00 by slow addition of 10M NaOH solution. 15 mM, 3 mM, 1 mM, 50 μM, and 50 nM EDTA solutions were prepared by dilution of the starting 150 mM solution of EDTA with Milli-Q water, and pH of the solution was measured before each measurement.

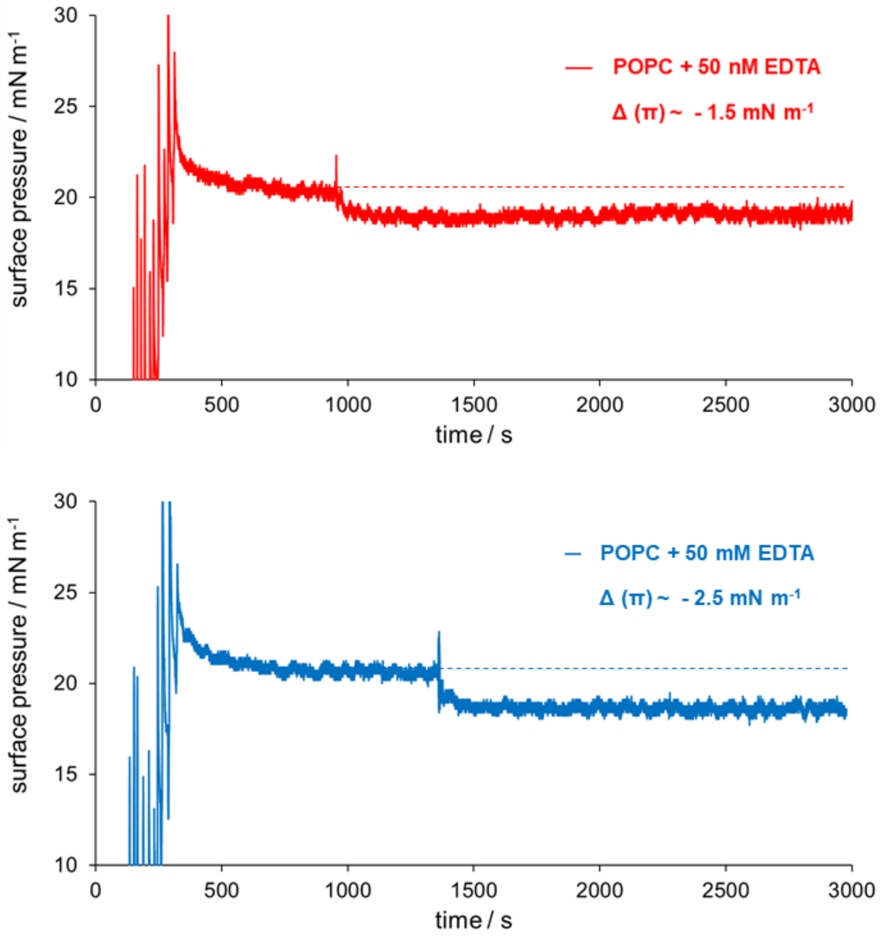


**Figure S1**. Kinetics of POPC monolayer surface pressure changes before and after addition of EDTA using cone Langmuir trough setup with NIMA sensor. The final concentration in the solution is 50 nM at pH = 5.5 (blue, upper panel) and 50 $\mu$M at pH = 6.4 (red, bottom panel).

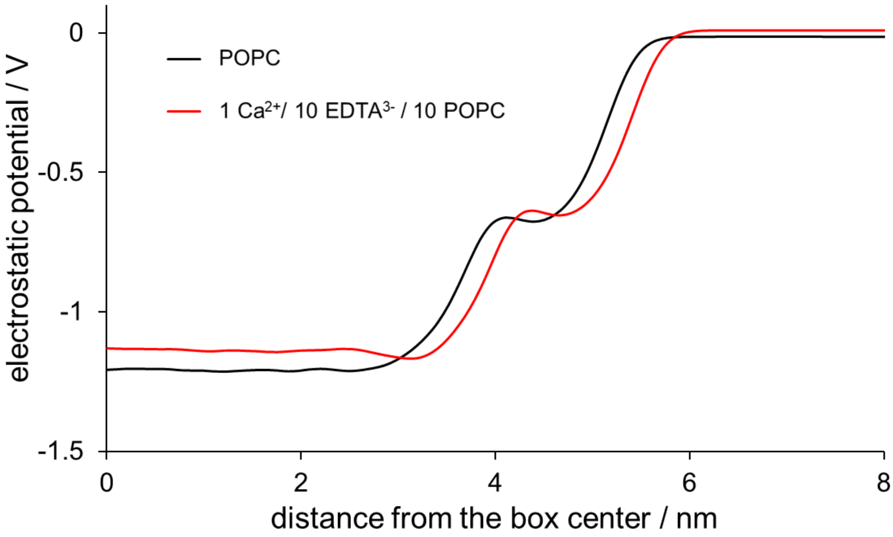


**Figure S2.** Electrostatic potential of the reference POPC (black line) and 1 $Ca^{2+}$ /10 $EDTA^{3-}$ /10 POPC systems.

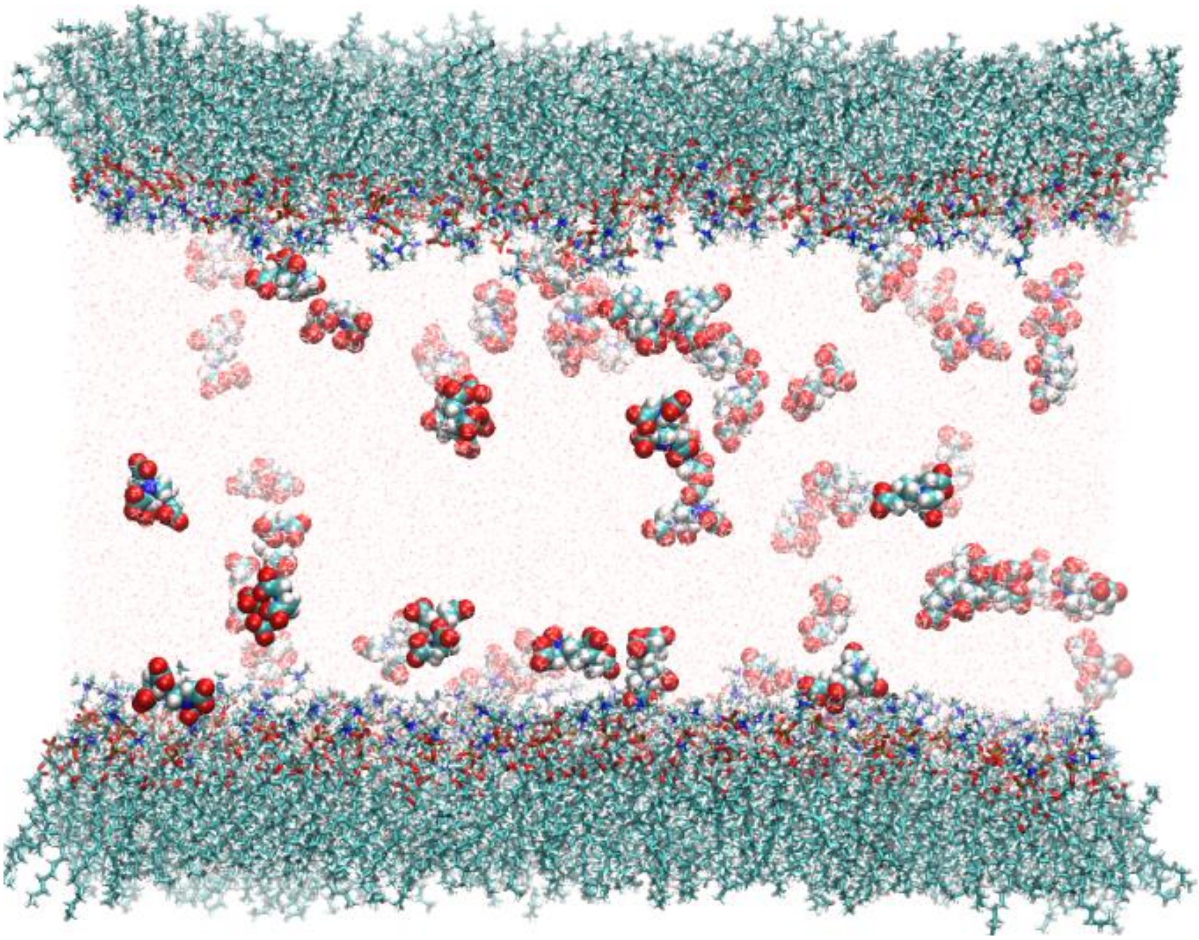

**Figure S3.** An example snapshot from MD simulations for 1 $Ca^{2+}$ /1 $EDTA^{3-}$ /10 POPC system.

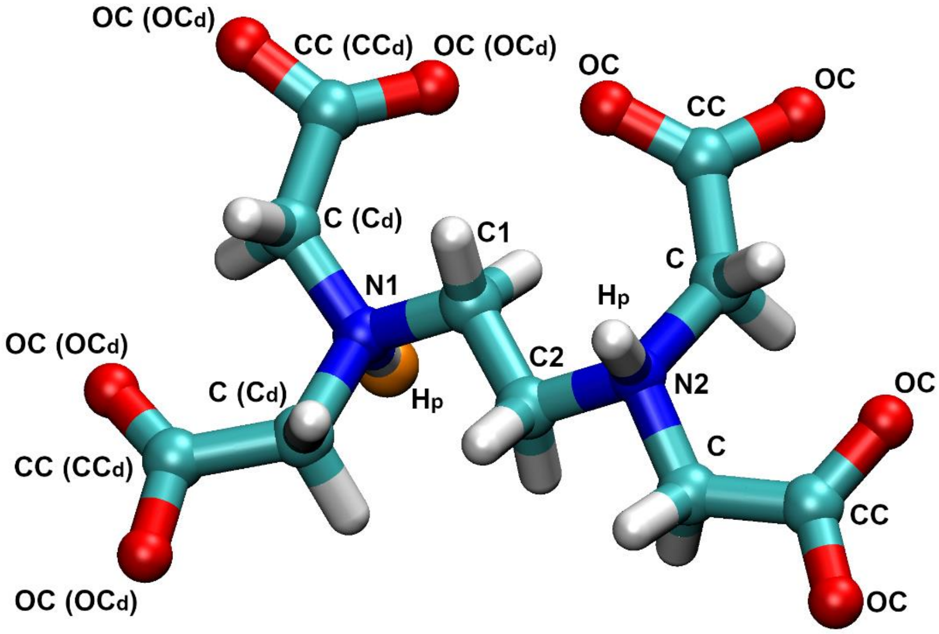


**Figure S4.** An MD model of EDTA anions. The structure of $EDTA^{2-}$ is shown, while $EDTA^{3-}$ is achieved by removing one of the protons shown as an enlarged orange sphere.

**Table S2.** Atomic partial charges for EDTA anions, see Figure S4 for atomic name assignment. All hydrogens that are not explicitly labeled correspond to atom name H. $C_d$, $CC_d$, and $OC_d$ atoms refer only to $EDTA^{3-}$ protonation form.

| Atom name | $EDTA^{2-}$ | $EDTA^{3-}$ |
|---|---|---|
| N1 | -0.134 | -0.716 |
| N2 | -0.134 | -0.148 |
| C1 | -0.05 | 0.108 |
| C2 | -0.05 | -0.034 |
| H | 0.09 | 0.09 |
| $H_p$ | 0.27 | 0.27 |
| C | -0.086 | -0.086 |
| CC | 0.603 | 0.603 |
| OC | -0.665 | -0.665 |
| $C_d$ | - | -0.011 |
| $CC_d$ | - | 0.564 |
| $OC_d$ | - | -0.76 |

### Supporting Information References
